# Micro-elastography of biopsies

**DOI:** 10.64898/2026.03.17.712283

**Authors:** Sibylle Gregoire, Bruno Giammarinaro, Domitille Le Quéré, Malaury Devissi, Axelle Brulport, Stefan Catheline

## Abstract

Micro-elastography is an optical technique that studies elastic waves for the mechanical characterisation of micrometric objects, such as cells. We propose to adapt this technique for the characterisation of millimetre-sized samples using a white light microscope. The objective is to perform a rapid, global characterisation of the elasticity of a biopsy. The millimetre-sized samples to be characterized are embedded in an agarose gel. A vibrator generates shear waves in this gel that transmit naturally inside the sample. This technique removes the need for precise manipulation of the wave source. A high-speed camera records the propagation of the waves in the sample. Their velocity is calculated using a noise correlation approach. Due to the lack of millimetric phantoms of calibrated elasticity, we choose to validate this method with a three step process. The experimental setup is first validated on homogeneous gels, then on biological samples of increasing elasticity, biopsies of beef liver hardened by heating, and finally on biological samples of clinical interest: biopsies of mouse endometrium. This method can be applied to all types of biological tissue, paving the way for rapid mechanical characterization of biopsies.

## Introduction

At the micrometric level, there is a wide range of techniques for estimating mechanical properties. Atomic force microscopy (AFM) sequentially scans the tissue to be characterized using a tip that applies a calibrated force. Cell monolayer rheometry measures the deformation of a cell sheet sheared by a commercial rheometer^1^. Parallel plate rheometry measures the elasticity of single cells held between two plates, one rigid and one flexible and oscillating^2^. Optical streching refers to the use of lasers to create an optical tweezers in which a single cell is stretched^1^. Magneto-cytometry, measure the mechanical response of a cell, functionalized beforehand with magnetic bead, when subjected to an electromagnetic field^2^. Static micro-elastography uses optical coherence tomography (OCT) to estimate the deformations of the object to be characterized when subjected to compression. The different layers of the skin^3^, breast cancer cells^4^ or, more recently, mouse ovaries^5^ have thus been studied. All these techniques are quasi-static techniques. They are sometimes local (static micro-elastography, AFM, magneto-cytometry), sometimes global (rheometry, optical streching). It is difficult to obtain quantitative results with these static techniques due to the difficulty of estimating the applied stress field. Wu et al^2^, compared different methods (AFM, optical streching, parallel plate rheometry, cell monolayer rheometry and magnetic cytometry) on the same breast cancer cell line and obtained high intermethod variability, with values ranging from 1 to 100-fold.

On the contrary, shear wave elastography is more easily quantitative and is also developed at this scale^6–8^. Shear wave elastography is a method for quantitatively estimating the elasticity of biological tissue. The propagation of shear waves naturally present in the tissue or induced by a vibration system is observed with an imaging system that can be ultrasound imaging^9^, magnetic resonance imaging^10^, optical imaging^11^ and more recently X-ray imaging^12^. The speed of shear waves is driven by the elasticity of the tissue. Historically, shear wave elastography was developed to measure the elasticity of whole organs or centimeter-sized samples. The development of optical elastography has led to dynamic micro-elastography, i.e the characterization of samples of micrometric size studying elastic waves propagation.

The smaller the sample, the shorter the wavelength required to characterize it. Therefore the higher the frequency of the shear wave field used. In biological tissues, high frequencies no longer propagate and become evanescent beyond a so-called cutoff frequency, which is approximately 20 kHz in soft tissues^13^. The propagation of high-frequency shear wave fields is observed in samples of small size like cell aggregates called spheroids measuring hundred of micrometers and individual cells measuring a few dozen micrometers^14,15^.

In this manuscript, we propose to adapt dynamic microelastography to the characterization of biopsies. The samples are millimeter-sized or smaller. The scale studied is therefore mesoscopic, halfway between conventional elastography and microelastography.

To reduce the complexity and cost of the system, we propose to use a white light transmission microscope instead of an interferometric method. This technique requires the sample to be transparent enough to allow visualisation through transmission. In this study, the elasticity of the sample is measured in about a hundred milliseconds. This rapid measurement limits the alteration of the physiological characteristics of the tissues. In fact, cell death is associated with cytoskeletal reorganization^16^ and changes in tissue mechanical properties^17^.

The experimental setup used is based on previous work^13,18^, and has been adapted to measure millimeter-sized tissue samples. The samples are embedded in an agarose gel. Waves are generated by a vibrator in contact with the gel, and are naturally transmitted to the tissue inside the gel. This technique doest not need precise micro-manipulation of the wave source.

The objective of this manuscript is to characterize and validate the ability of this experimental setup to measure the elasticity of biopsies. To our knowledge, there are no millimeter-scale equivalents to calibrated elasticity phantoms, such as the CIRS Module 049 for ultrasound elastography. We therefore chose another method. First, homogeneous agarose gels are studied. The elasticity of gels with increasing concentrations is measured. Biological tissue is then studied in order to validate the sensitivity of the technique to variations in elasticity. We chose to study the stiffening of beef liver by comparing the elasticity of samples cooked for between 0 and 5 minutes in a water bath. Finally, we are studying the elasticity of a heterogeneous tissue of clinical interest : the endometrium.

The endometrium is a biological tissue with a complex histology, containing stroma and glands, and composed of several cell types (mainly epithelial and stromal cells). It physiologically lines the inner wall of the uterus and allows embryo implantation. Due to the menstrual cycle, the tissue undergoes remodeling and angiogenesis, which affects its elasticity^19^. The elasticity of the endometrium is also studied to assess endometrial receptivity and predict pregnancy outcome. Using shear wave elastography, Zhang et al^20^, found higher endometrial elasticity in pregnant women after embryo implantation^20^. The main pathologies affecting the endometrium are endometriosis, polyps, synechiae, endometritis, and cancer. All of these pathologies are associated with inflammation that causes fibrosis of the tissue and alters its elasticity. Women with endometrial carcinoma have a stiffer endometrium than women in the control group^21^. The elasticity of the endometrium also increases with the number of voluntary abortions^22^. The study of endometrial elasticity is therefore of interest for understanding the physiology and pathophysiology of this tissue.

## Experimental

### Micro-elastography setup

Figure 1 (A) describes the experimental setup for characterizing biopsies. An ultra-fast camera records the propagation of shear waves in a millimeter-sized tissue sample. The sample is embedded in an agarose gel, the waves are generated in the gel with a piezoelectric actuator, and then are transmitted from the gel to the sample. An example of millimeter-sized tissue is photographed in Figure 1 (B). It is contained in a 1% agarose gel in a Petri dish. The Petri dish is filled with water, and a thin layer of glass is placed in contact with the surface of the fluid. The water and glass prevent the shadow of the waves on the surface of the gel from being visible when deeper layers are probed. These surface waves can be studied to characterize the agarose gel, but they constitute measurement noise for the waves of interest propagating in the sample.

**Fig. 1.**
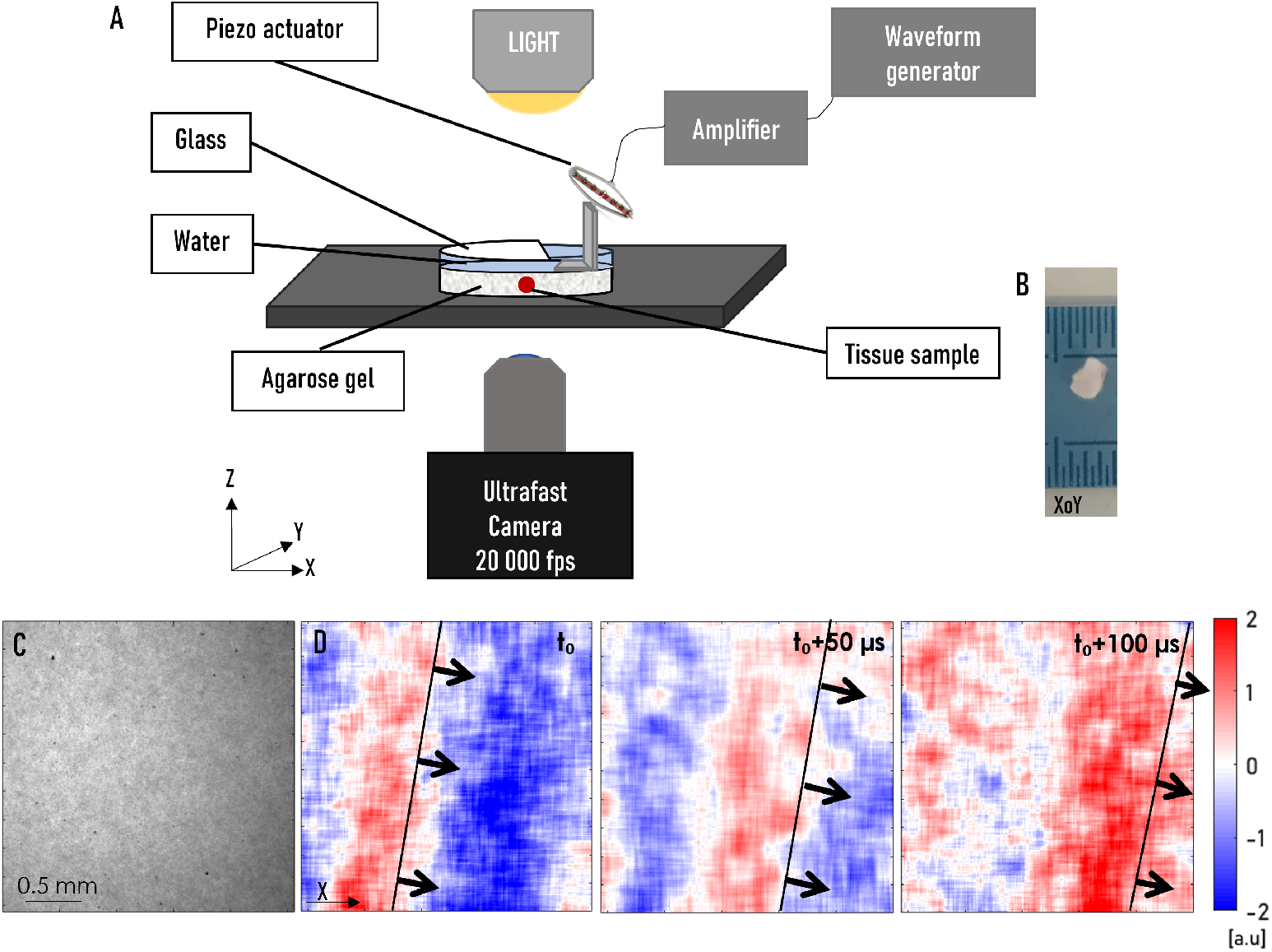
(A) Experimental setup for optical elastography for biopsy characterization. An ultra-fast camera records the propagation of shear waves in a millimeter-sized tissue sample. These waves are generated by a piezoelectric actuator in an agarose gel that serves as a wave transducer. (B) Photograph of a tissue sample in agarose gel. (C) Microscopic image of a homogeneous agarose gel sample (D) Displacement field in this gel at an arbitrary time *t*_0_, then at *t*_0_ + 50 *µs* and *t*_0_ + 100 *µs*

An ultra-fast camera (Phantom v12.1) is mounted on an inverted microscope (Nikon), equipped with a 4x or 10x magnification lens. The field of view is 512*512 pixels, acquired at a rate of 20,000 frames per second. A piezoelectric actuator (Cedrat Technologies, APA100M vibrator) mounted with a 3D-printed Lshaped adapter is placed in contact with the gel surface. The signal sent to the actuator is a sinusoidal signal ranging from 2.5 to 8 kHz with a duration of 100 ms. An amplifier (Cedrat Technologies, LC75B, LA75A) adapted to the piezoelectric actuator is used. The acquisition time is 100 ms. 2000 images are recorded per acquisition.

Homogeneous agarose gels at different concentrations are studied using the experimental setup described above, but without tissue samples. Agarose (VWR) at 0.5% w/v, 1% w/v, 1.5% w/v, and 2% w/v is dissolved by heating under constant stirring. An aqueous solution (0.05 w/v) of aluminium oxide microparticles (1 micron, Logitech) is added before the solution is poured into Petri dishes. To evaluate reproducibility, three gels are made for each concentration. Figure 1 (A) is a 4x enlarged image of a 1% agarose gel ; the visible dots are the incorporated microparticles.

### Estimation of the displacement field

In order to visualize the movements in the sample, the displacements in the image between two successive images are calculated. To do this, the phase of the displacement signal of each point in the image, is estimated in a given direction, by applying the Hilbert transform in the direction considered. The phases obtained are then subtracted in pairs to obtain the displacement field^23^. The energy of the displacement signal is then calculated over time using a sliding window. To limit the noise level, displacement signals on windows with energy less than one-third of the maximum energy obtained are set to zero. This operation reduces the noise level by considering only the most significant displacements. Figure 1 (B) shows the displacement field for three consecutive instants. The passage of a wave front through the gel can be observed.

### Estimation of shear wave velocity

To estimate the speed of the observed waves, we use a noise correlation method derived from seismology^24,25^. The main idea is that the time-reversed (TR) field of a diffuse isotropic wave field, *ϕ*, calculated between two receivers separated by a distance *r*, is directly related to the Green’s function between them. The normalized TR field, *ϕ* ^*TR*^(*r, t*), is defined in the time domain by equation 1, where ⊛ represents the time convolution product and 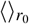 is a spatial average taken at any arbitrary point.

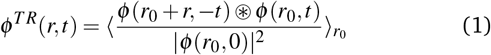

In the case of a diffuse field, its Fourier transform 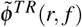 is related to the imaginary part of the Green’s function^26^ by equation 2, where f is the frequency.

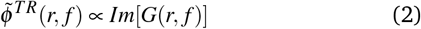

The Green’s function of a 3D diffuse field in a soft, isotropic, and homogeneous solid is given by equation 3, which is shown below, where *j*_0_ and *j*_2_ are the spherical Bessel functions of order zero and two, *c*_*s*_ is the shear wave velocity, and *θ* is the angle between the direction of displacement and the polarization of the source.

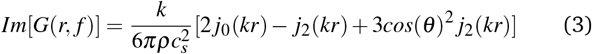

We assume that in our experiment we generate a diffuse field and that experimentally the information in all directions of the imaging plane are collected. An integration over all directions *θ* is performed.

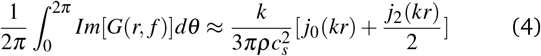

Furthermore, for small distance *r* compared to the wavelength, the term *j*_0_ dominates over the term 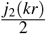. We can therefore approximate *Im*[*G*(*r, f*_*c*_)] using equation 5, where *f*_*c*_ is the central frequency of the field, *λ*_*c*_ is its wavelength, and *c*_*s*_ is the shear wave velocity.

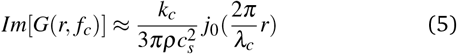

Experimentally, the TR field at time zero *ϕ* ^*TR*^(*r*, 0), also called the focal spot, is calculated using equation 1 by averaging the points *r*_0_ describing a 10:10 grid centered on the central pixel. Given the width of the field’s frequency band, we estimate that 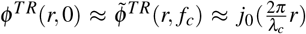. A nonlinear regression using a function *j*_0_ is applied to determine the wavelength. Figure 2 shows the results obtained on a 1% agarose gel sample. Figure 2 (A) shows the focal spot *ϕ* ^*TR*^(*r*, 0), in black, on an arbitrary space line. The width of the focal spot is estimated by the wavelength of the fit of the function *j*_0_, plotted in red.

**Fig. 2.**
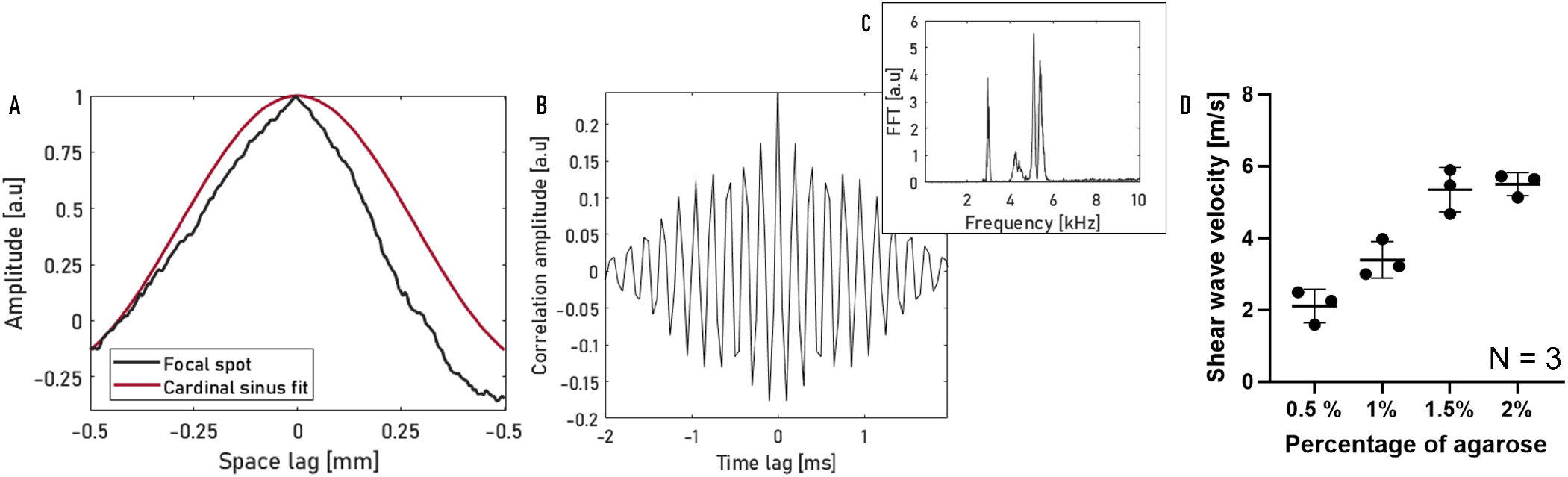
(A) Focal spot *ϕ* ^*TR*^(*r*, 0) at the center point of the field of view and its estimation by a cardinal sine in red for an acquisition performed on a homogeneous agarose gel. (B) Autocorrelation in time of the displacement field *ϕ* ^*TR*^(0, *t*) and (C) its spectrum. (D) Shear wave velocity in homogeneous agarose gel samples with increasing agarose concentration. The mean value and standard deviation are calculated from the three replicates of gels made at each concentration.

In order to estimate the center frequency of the displacement field, the autocorrelation of the signal at each point, *ϕ* ^*TR*^(0, *t*), is calculated. An example is given for a homogeneous gel in Figure 2 (B). The Fourier transform of *ϕ* ^*TR*^(0, *t*) is shown in Figure 2 (C). The spectrum is centered around 5 kHz. Although the control signal is between 2.5 and 8 kHz, the spectrum only shows the signal between 3 and 6 kHz. This is due, on the one hand, to the response of the vibrator and the L-shaped adapter attached to it and, on the other hand, to the viscoelastic properties of the gel. The central frequency is estimated to be the frequency with the highest energy in the spectrum. In the example Figure 2 (C), it is 5.1 kHz.

## Results

### Characterization on homogeneous agarose gels

The shear wave velocity obtained from spatial and temporal focal spots on acquisitions on homogeneous agarose gels is shown in Figure 2 (D). Each point corresponds to the average shear wave velocity obtained from five acquisitions performed by varying the area probed on a gel. As expected, an increase in shear wave velocity is observed as the agarose gel concentration increases: *c*_0.5%_ = 2.1 ± 0.5 m/s, *c*_1%_ = 3.4 ± 0.5 m/s, *c*_1.5%_ = 5.3 ± 0.6 m/s, *c*_2%_ = 5.5 ± 0.3 m/s. This represents a 160% increase in the average shear wave velocity in a 2% agarose gel compared to a 0.5% agarose gel. This experiment therefore validates our approach. This micro-elatography setup is able to probe elasticity.

### Monitoring the elasticity of beef liver during heating in a water bath

The next step is to apply our method to genuine biological tissues. We are looking to observe variations in elasticity. Variations in the elasticity of biological tissues when heated are studied as part of real-time monitoring of high-intensity focused ultrasound (HIFU) treatments^27–29^. Muscles exhibit complex behavior when the temperature rises. During heating, they first soften before hardening^28^. The liver exhibits simpler behavior. Sapin-de Brosses et al^28^ showed hardening above 45°C, Arnal et al^27^ showed a 20% to 30% increase in shear modulus after HIFU treatment, and Barrère et al^30^ showed an increase in shear wave velocity from 1 to m/s for an increase of 30°C, using the noise correlation methods described earlier. We therefore chose to study the hardening of the liver subjected to increasing heating times.

The samples are prepared as described in Figure 3 (A). Samples of beef liver measuring a few centimeters are placed in tubes (Falcon 15 mL) and immersed in boiling water for 0 to 5 minutes. From these samples measuring a few centimeters, samples measuring 1 to 2 mm^3^ are cut using a scalpel. These samples, which are typically the size of a biopsy, are then placed in Petri dishes filled with 1% agarose gel at 37°C. In order to get an idea of reproducibility, the heating process is repeated for three samples.

**Fig. 3.**
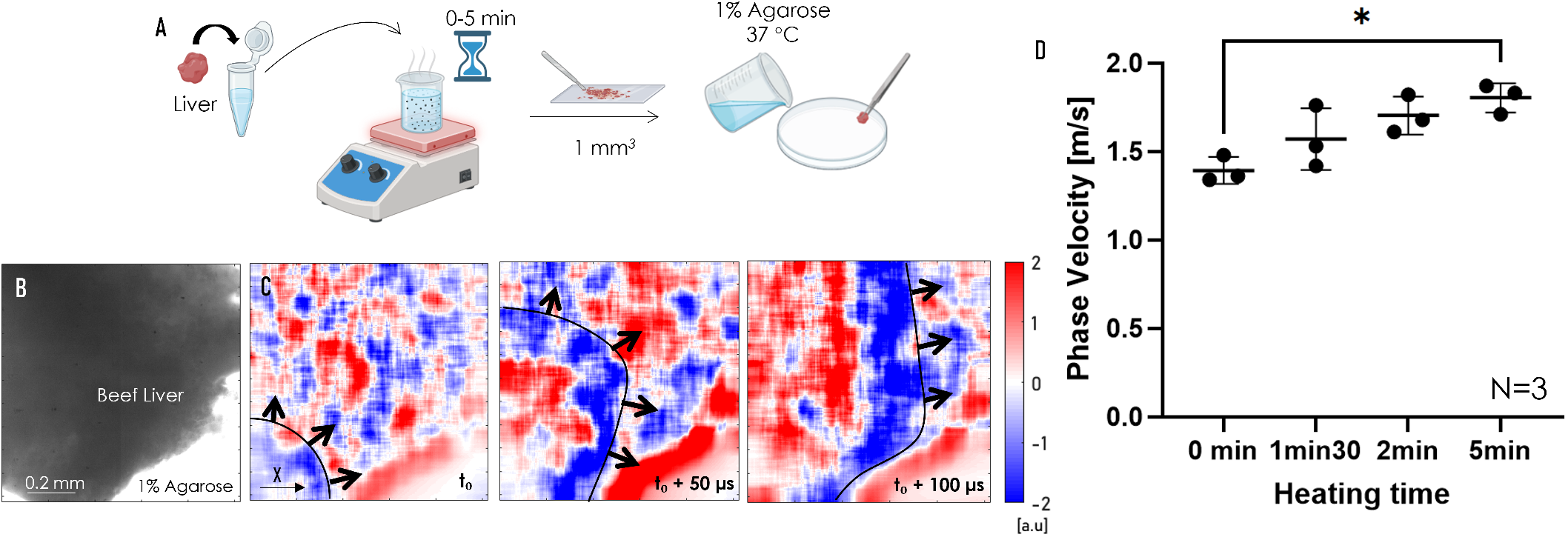
(A) Diagram of the different stages of liver sample preparation. Created with Biorender. (B) Microscopic image of a beef liver sample embedded in 1% agarose gel. (C) Displacement field in the sample at an arbitrary time *t*_0_, then at *t*_0_ + 50 *µs* and *t*_0_ + 100 *µs*. The black arrows indicate the direction of the wave fronts. (D) Shear wave velocity in beef liver cooked for 0 min, 1 min 30 s, 2 min, and 5 min. The mean value and standard deviation for the three experiments are shown.

Figure 3 (B) is a 10x magnified image of a liver sample embedded in agarose. Although the liver appears opaque and homogeneous, we can see some roughness. These granularities allow for good efficiency of the tracking algorithm based on phase difference measurements. Figure 3 (C) shows the displacement field in the X direction in this sample for three consecutive moments.

In order to estimate the group velocity of the observed waves, we use the noise correlation method described earlier. On one side, the focal spot *ϕ* ^*TR*^(*r*, 0) is computed to estimate the average wavelength, and on the other side, the temporal autocorrelation *ϕ* ^*TR*^(0, *t*) is computed to estimate the central frequency. The results are shown in Figure 3 (D). The mean value and standard deviation for the three experiments are displayed. The liver has an initial shear wave velocity of *c*_0*min*_ = 1.4 ± 0.1 m/s. An increase in shear wave velocity is observed with time spent in the water bath. After 1 minute 30 seconds in boiling water, the liver has a shear wave velocity of *c*_1*min*30_ = 1.6 ± 0.2 m/s, after 2 minutes *c*_2*min*_ = 1.7 ± 0.1 m/s, and after 5 minutes, *c*_5*min*_ = 1.8 ± 0.1 m/s, representing a 30% increase from the initial value. A nonparametric Kruskal-Wallis multiple comparison statistical test was performed using GraphPad Prism 10 software. A significant difference at the 5% threshold was observed between the shear wave velocities of the 0 and 5 minute heated groups (*p* = 0.04).

### Elastography of a complex tissue: the murine endometrium

We are now seeking to characterize a heterogeneous tissue whose elasticity is of clinical interest: the endometrium. As the mechanical properties of the endometrium are unknown to us, we choose to compare it to the liver, a more homogeneous tissue that has been studied extensively in elastography. The aim is to obtain an initial estimate of endometrial elasticity. This study is too preliminary to justify collecting samples from patients during an invasive examination. We have therefore chosen to work with animal models. The mouse was chosen because it is a widely available model that allows for the simple and rapid collection of both liver and endometrial samples from an individual. The number of individuals was limited to five in order to achieve adequate statistical power without resorting to an excessive number of individuals. The protocol for preparing samples is illustrated in the Figure 4 (A). The uterus and liver of five individuals were removed by dissection. Between dissection and characterization, the tissues are stored on ice. A 1% agarose gel is prepared and transferred to a Petri dish once it has cooled to 37 degrees. The samples are then added to the gel before it solidifies.

**Fig. 4.**
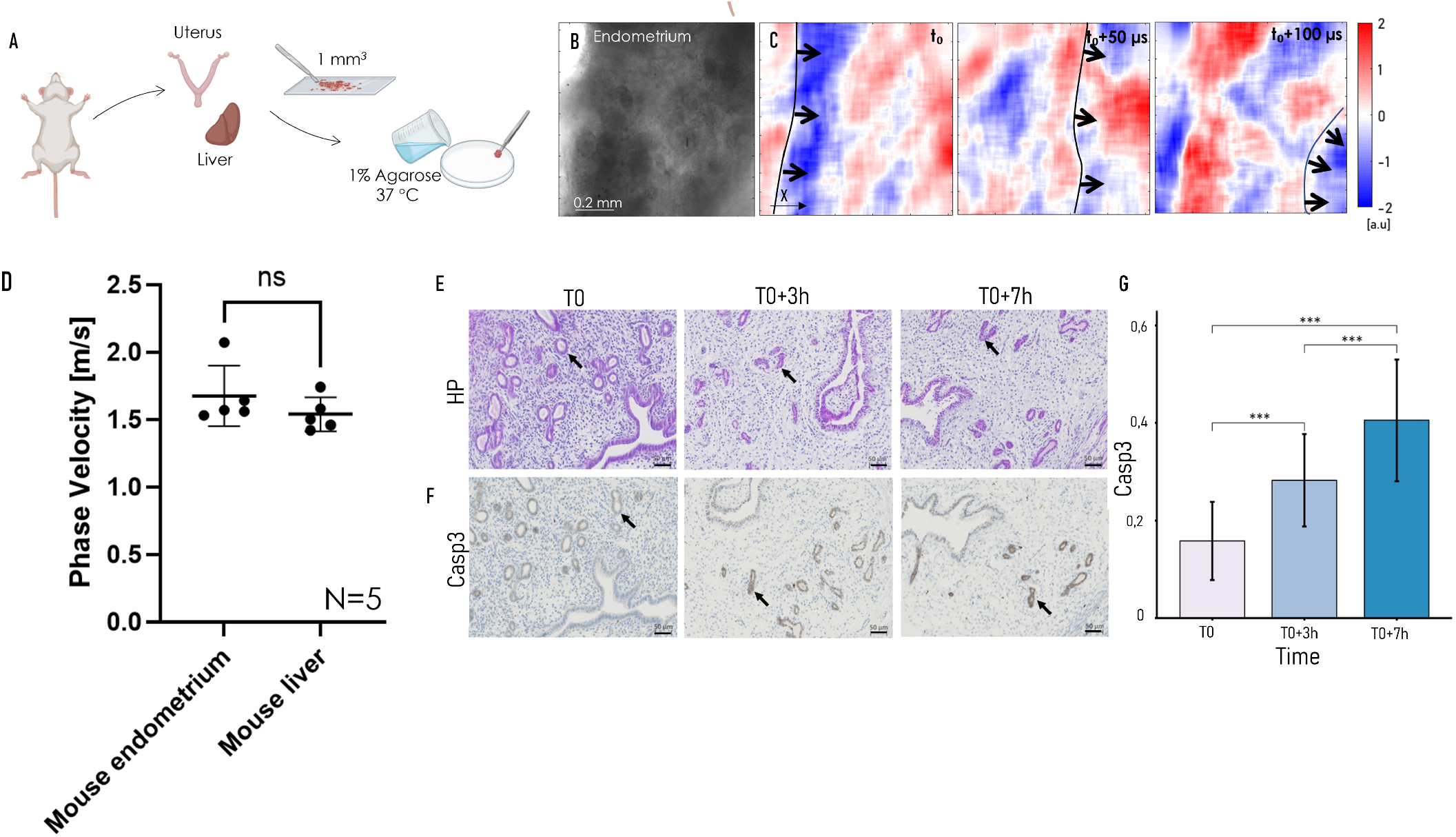
(A) Scheme of the different steps in the preparation of mouse endometrial and liver samples. Created with Biorender. (B) Microscopic image of an endometrial liver sample in 1% agarose gel. (C) Displacement field in the sample at an arbitrary time *t*_0_, then at *t*_0_ + 50 *µs* and *t*_0_ + 100 *µs*. The black arrows indicate the direction of the wave fronts. (D) Shear wave velocity in mouse liver and endometrium. The mean value and standard deviation for five individuals are shown. Monitoring of tissue physiological properties over time in 1% agarose gel. Evaluation of tissue structure on hematoxylin and eosin-stained sections (E), monitoring of cell viability on sections labeled with anti-caspase 3 antibody (F), and relative quantification of caspase 3 labeling (G). Black arrows highlight the reduction in glandular volume.

Figure 4 (B) is a 10x enlarged image of an endometrial sample embedded in agarose. It shows the complex structure of the endometrium. The transparency varies across different areas of the image, indicating a difference in thickness within the sample. Figure 4 (C) shows the displacement field in the X direction in this sample for three consecutive time points. The wave fronts are clearly visible, their movements highlighted by black arrows. These wave fronts are not flat due to the internal structures of the tissue. The images acquired in the murine liver are similar to those acquired in the bovine liver in Figures 3 (C,D). They are not shown here for the sake of conciseness.

To estimate elasticity, the noise correlation method is applied. The results are shown in Figure 4 (D). The endometrium appears to be slightly stiffer than the liver. A nonparametric Mann-Whitney statistical test is performed and no significant difference is found between the two groups. The endometrium and liver in mice seems to have comparable elasticity.

### Estimation of tissue viability after incorporation into agarose

Cell death is associated with a change in the mechanical properties of tissues^16,17^. We seek to determine whether the mechanical properties of biopsies are altered at the time of measurement in agarose compared to the time of sampling. To do this, we check the viability of the samples after 5 minutes (T0), 3 hours (T0 + 3h) and 7 hours (T0 + 7h) in an agarose gel. First, the structure of the tissue and its state of dehydration are revealed by studying histological sections stained with haematoxylin and eosin. Secondly, sections marked with a caspase-3 antibody, which is an indicator of cell viability, are examined.

Endometrial biopsies were taken from the mice immediately after sacrifice and embedded in 1% agarose at 37°C. The biopsies were removed from the agarose after 5 minutes (T0), 3 hours (T0 + 3h), and 7 hours (T0 + 7h) at room temperature. After removing the biopsies from the agarose, the tissues were fixed for 1 hour in formalin, dehydrated, and embedded in paraffin. Three-micrometer sections were stained with hematoxylin and eosin and immunohistochemically stained for caspase 3 (Cell Signaling, No. 9664) using the following parameters (dilution: 1/400, incubation: 1 hour at 37°C) with the assistance of the “Lyon-Est Quantitative Imaging Center” platform at the University of Lyon I. Antigen retrieval was performed with Tris EDTA buffer (pH = 8, 40 min, 95°C). Detection was performed with secondary antibodies conjugated to streptavidin-HRP, and the sections were counterstained with hematoxylin.

The photos shown in Figure 4 (E,F) were taken using the AXIO SCAN 7 slide scanner at 10x magnification. Figure 4 (E), showing sections stained with hematoxylin and eosin, highlights the tissue structure. Dehydration of the endometrial samples is observed over time in agarose. This is evidenced by a reduction in glandular volume, highlighted with black arrows in Figure 4 (E). This dehydration phenomenon is associated with a decrease in cell viability.

Caspase-3 staining is a marker of apoptosis. The increase in cell death is thus associated with a significant intensification of caspase-3 staining. Figure 4 (F) an increase in the staining can be observed by eye in glandular epithelial cells, highlighted with black arrows. Using MatLab software, the cell area of 10 glands per slide was manually counted. The density of brown-stained pixels was then evaluated using a threshold (red [40, 120], green [35, 115], blue [30, 110]) on the selected region. The staining was quantified as follows: brown area/total gland area. This quantification was performed on three biological replicates for each time point. To compare the density of brown staining at T0, T0 + 3h, and T0 + 7h, a paired t-test with Bonferroni correction was performed using R 4.4.2. Comparison between T0 and T0 + 3h reveals a 1.75-fold increase in caspase-3 labeling with a corresponding p-value of 1.31E-5. Similarly, the comparison between T0 and T0 + 7h showed a 2.56-fold increase with a p-value of 3.34E-15, see Figure 4 (G).

The intensity of caspase-3 staining at T0 is similar to that observed in endometrial biopsies from healthy women^31^. The inclusion of tissue in agarose has no immediate impact on tissue viability and structure. However, the increase in the intensity of this apoptosis marker and tissue dehydration at T0 + 3h and T0 + 7h shows the importance of performing acquisitions quickly.

## Discussion

Inspired by previous work on optical elastography^13,15,18^, micro-elastography is suitable for measuring the elasticity of millimeter-sized samples. A layer of water and a thin layer of glass are added to prevent deep surface wave shadows. The samples are immersed in a gel. The wave source is indirect: the piezoelectric actuator generates waves in the agarose gel, which then propagate into the sample to be characterized. This facilitates the generation of a wave field in the sample, eliminating the need for micro-manipulation of the source.

The technique is validated in three stages. First, the elasticity of agarose gels of increasing concentration is characterized. A 160% increase in shear wave velocity is observed when comparing a 0.5% agarose gel with a 2% agarose gel. The next validation step is to estimate the elasticity of real biological tissue samples. We choose beef liver, a homogeneous biological tissue studied in the literature. The values reported for this tissue are 1 to 2 m/s at a few hundred Hz^30,32,33^. A similar order of magnitude is obtained: *c*_0*min*_ = 1.4 ± 0.1m/s. The elasticity of beef liver cooked in a water bath for 0 to 5 minutes is then measured. As predicted by the literature^27,28,30^, an increase in shear wave velocity is observed. In particular, the liver is significantly stiffer after 5 minutes in boiling water, with an increase of 30% (0.04, P < 0.05).

Finally, the elasticity of biopsies from clinically relevant heterogeneous tissue, the endometrium, is studied and compared to that of a known tissue, mouse liver. The viability of endometrial samples at the time of measurement is verified. The need for rapid acquisition is highlighted by a significant increase in cell death and tissue dehydration within 3 hours. The shear wave velocity measured in mouse liver samples is *c*_*liver*_ = 1.5 ± 0.1 m/s. The shear wave velocity measured in mouse endometrial samples is slightly higher *c*_*endo*_ = 1.7 ± 0.2 m/s. However, there is no significant difference between the shear velocities obtained in the two tissues.

Under the assumptions of a homogeneous, isotropic, infinite medium, approximate estimate of Young’s modulus are derived, *E*_*liver*_ ≈ 3*ρc*^2^ = 7 ± 1 kPa and *E*_*endo*_ = 8 ± 2 kPa.. The dimensions of the sample are millimeters. We can assume that for certain samples there are guiding effects, where the assumption of an infinite medium is no longer valid. In addition, the endometrium is a heterogeneous tissue (see the histological sections shown in Figure 4 (E)). The above estimate of elasticity is therefore a first approximation that facilitates comparison of the microelastography estimate with the literature. To our knowledge, no values have been published concerning the elasticity of mouse endometrium, we are therefore comparing our elasticity estimation to human endometrium. Jiang et al^34^, using MRI elastography, found Young’s moduli of 6 to 9 kPa for frequencies of several tens of Hertz. Jiao et al^22^ studied the endometrium using ultrasound elastography and found a magnitude of approximately 10 ± 4 *kPa* at several hundred Hertz. The same order of magnitude of elasticity is obtanied in this manuscript. However, these studies estimate elasticity at frequencies well below those used in our work (2.5 to 8 kHz), in human endometrium, in vivo. It is therefore difficult to draw conclusions, given the difficulty of comparing measurements at a few Hz with measurements at a few kHz, in different species, in vivo and ex vivo.

Jiang et al^34^ observed a 30% variation in shear modulus due to the menstrual cycle. In mice, the endometrium remodels throughout the individual’s cycle, which is not the case for the liver. Inter-individual variability in endometrial samples was found to be slightly higher than in liver samples. These results were expected, as the period of the mouse cycle at the time of measurement was random.

There are several areas for improvement in the technique of white light micro-elastography of biopsies. First, it can be assumed that the geometry of the biopsies on which the acquisitions are performed has an impact, as the high frequencies generated in the biopsy can be more or less guided. In this study, a scalpel was used to cut the samples, but in clinical practice, curettes or biopsy guns are used to take biopsies in vivo. One possible improvement would be to use a biopsy gun, which produces biopsies with calibrated geometries. The shape of the wave field may also influence the measurement. Indeed, by adjusting the Bessel function *j*_0_ to the experimental focal spots, we assume that the observed field is a diffuse 3D field. This assumption is not verified in the case of plane waves in a single direction, where a sinusoidal shape is expected rather than a cardinal sine shape. This leads to an overestimation of the wave velocity. This can be corrected by adapting the fitting function to the wave field generated in the sample.

## Conclusions

In conclusion, we developed an experimental setup to characterize the elasticity of biopsies using a white light microscope and a high-speed camera. High-frequency waves are generated in an agarose gel and transmitted to the biopsy included in this gel. The wavelengths and central frequencies of the shear wave field generated in the biopsy are measured by noise correlation. We validated the micro-elastography technique for biopsies in three stages. First, the velocities of homogeneous gels with increasing elasticity are measured. Next, variations in the elasticity of biological tissue are measured: the hardening of beef liver during cooking. Finally, a heterogeneous biological tissue, mouse endometrium, is studied and found to have elasticity comparable to that of mouse liver: *E*_*liver*_ = 7 ± 1 kPa, *E*_*endo*_ = 8 ± 2 kPa. White light micro-elastography can be used on all types of biological tissue that are sufficiently optically transparent, paving the way for rapid mechanical characterization of all types of biopsies. In particular, a clinical trial collecting endometrial biopsies from women with endometriosis has been initiated to investigate elasticity as a biomarker of fibrosis in women with endometriosis.

## Author contributions

### Data availability

Data supporting the findings of this study are available from the corresponding author on a reasonable request.

## Acknowledgements

We thank Inas H Faris and Jose Cortes for their assistance in designing the setup, particularly in adding water and glass to eliminate optical artifacts. We also thank Marine Simonneau for her help in dissecting the mice.

